# Dose- and outcome-dependent effects of bacterial infection on female fecundity in *Drosophila melanogaster*

**DOI:** 10.1101/2025.11.22.689874

**Authors:** Aabeer Basu, Kimaya Tekade, Nagaraj Guru Prasad

## Abstract

The virulence of a pathogenic infection challenge can manifest in the form of increased host mortality rates and/or reduced host fecundity. Reduced fecundity in infected hosts can result from resource-allocation trade-offs: increased investment in immune defences depletes the common pool of resources, which are also required for reproduction. Alternatively, reduced fecundity may result from damage to host organs, especially reproductive organs, caused by the infection. We infected *Drosophila melanogaster* females with the bacterial pathogen *Enterococcus faecalis* at different infection doses and found that increasing the infection dose led to greater suppression of fecundity, without a corresponding increase in mortality rates. We further found that the reduction in host fecundity is contingent on the infection outcome (whether the host lives or dies after being infected), particularly in flies infected with a high dose of bacteria. Interestingly, survivors of the infection challenge had exhibited comparable fecundity irrespective of the dose used to infect them, whereas amongst females that died after infection, a higher infection dose led to a lower fecundity. We therefore propose that our results indicate that fecundity suppression in *E. faecalis*-infected females is likely caused by host organ damage rather than by diversion of limited resources towards immune function.

## Introduction

A central tenet in eco-immunology is that the defence against pathogens and parasites is physiologically costly for the host organism (Sheldon & Verhulst, 1996, Rolff & Siva-Jothy, 2003, Schulenburg et al., 2009, Sadd and Schmid-Hempel 2009). These defence costs can be constitutive or inducible, that is, the costs may manifest either in the absence or in the presence of an infection, respectively (Armitage et al., 2003, Schmid-Hempel 2005). The inducible costs are resource reallocation costs: when infected, the host invests excess resources in immune function, thereby depriving other critical physiological functions of essential resources (Lochmiller and Deerenberg 2000). Such resource reallocation costs commonly manifest as compromised life-history traits, for example, reduced reproductive output (Schmid-Hempel 2005). However, although resource reallocation from reproduction towards immune function can manifest as reduced host fecundity, a post-infection compromise in host fecundity does not necessarily imply a resource-based trade-off.

A pathogenic infection, in addition to resource trade-offs, can compromise host fecundity via damage to the host soma, especially the host reproductive tissue (Hurd 2001, Brandt and Schneider 2007). This damage can come from either the actions of the pathogen or those of the host’s own immune response (Brandt and Schneider 2007, Sadd and Siva-Jothy 2006). Therefore, it is difficult to attribute a post-infection reduction in host fecundity to a specific cause. Further complications arise when a pathogenic infection has no effect, or even increases, host fecundity (Minchella and Loverde 1981, Parker et al., 2011).

The observed and empirically measurable effects of a pathogenic infection on host fecundity are proposed to be contingent on various factors, including the host environment and physiology, and the biology of the pathogen (Sandland and Minchella 2003, Duffield et al., 2017, Nystrand and Dowling 2020). In this study, we explored how the infection dose (number of pathogen units introduced into the host body) and the outcome of infection (whether the host dies or survives by the end of the acute phase of infection) affect fecundity in *Drosophila melanogaster* females, both independently and interactively, when infected with *Enterococcus faecalis*, a known bacterial pathogen of this species (Lazzaro et al., 2006, Shirasu-Hiza and Schneider 2007, Dionne and Schneider 2008). Previous research in *D. melanogaster* has established that the effects of bacterial infection on female fecundity can depend on a variety of factors, including but not limited to pathogen identity and pathogenicity, host genotype and diet, and infection parameters such as the route and phase of infection (Brandt and Schneider 2007, McKean et al., 2008, Linder and Promislow 2009, Ye et al., 2009, Howick and Lazzaro 2014, Kutzer and Armitage 2016, Kutzer et al., 2018, Kutzer et al., 2019, Hudson et al., 2020, Basu et al., 2024, Adhikari et al., 2025).

In our experiments, we measured the survival, fecundity, and bacterial load upon death (BLUD) of individual females, infected at two different infection doses, with a two-fold difference in the number of bacteria introduced into the host. We find that infection challenge suppresses fecundity, with the degree of suppression being determined interactively by infection dose and outcome. However, infection-induced fecundity suppression is independent of infection-related mortality risk. We speculate that infection-induced damage to host soma drives fecundity suppression, assuming a higher infection dose causes greater damage, an assertion supported by other existing studies (Brandt and Schneider 2007, Sehgal et al., 2025).

## Materials and methods

### Pathogen handling and infection protocol

The Gram-positive bacterium *Enterococcus faecalis* (Lazzaro et al., 2006) was used in the experiments reported here. The bacteria are preserved as glycerol stock at −80 ^O^C. To obtain live bacterial cells for infections, 10 mL of lysogeny broth (Luria-Bertani Broth, Miler, HiMedia) is inoculated with the bacterial stock and incubated overnight with aeration (150 rpm shaker incubator) at 37 °C. 100 microliters of this primary culture are inoculated into 10 mL of fresh lysogeny broth and incubated for the necessary amount of time to obtain confluent cultures (OD600 = 1.0-1.2). The bacterial cells are then pelleted down using centrifugation and resuspended in sterile MgSO_4_ (10 mM) buffer at an optical density (OD_600_) of 1.0; OD_600_ = 1.0 for *E. faecalis* corresponds to 10^7^ cells/mL. Flies are infected, under light CO_2_ anaesthesia, by pricking the dorsolateral side of the thorax with a 0.1 mm Minutien pin (Fine Scientific Tools, CA, USA) dipped in the bacterial suspension. Sham-infections (controls for injury) are carried out in the same fashion, except by dipping the pins in sterile MgSO_4_ (10 mM) buffer.

### Host population

Flies from the <u>B</u>lue <u>R</u>idge <u>B</u>aseline <u>2</u> (BRB2) population – a large, lab-adapted, outbred *Drosophila melanogaster* population – were used as the focal flies in the experiments reported here. This population was established by hybridising 19 iso-female lines, which were founded from wild-caught females (Singh et al., 2015). The BRB2 population is maintained as a large, outbred population with a generation time of 14 days, comprising approximately 2,800 adults in each generation. Every generation, eggs are collected from the population cage (plexiglass cage: 25 cm length × 20 cm width × 15 cm height) and dispensed into vials (25 mm diameter × 90 mm height) with 8 mL banana-jaggery-yeast food medium at a density of 70 eggs per vial. 40 such vials are set up; the day of egg collection is demarcated as day 1. These vials are incubated at 25 °C, 50–60% RH, 12:12 hour LD cycle; under these conditions, the egg-to-adult development time for these flies is about 9–10 days. On day 12 post egg collection, all adults are transferred to a population cage and provided with fresh food plates (banana-jaggery-yeast food medium in a 90 mm Petri plate) supplemented with *ad libitum* live yeast paste. On day 14, a fresh food plate is provided in the cage, and 18 hours later, eggs are collected from this plate to initiate the next generation.

For each replicate of the experiments, eggs were collected from the BRB2 population cage and distributed into food vials containing 8 mL of standard food medium at a density of 70 eggs per vial. These vials were incubated under standard maintenance conditions. 12 days post egg-laying, adult flies were transferred to fresh food vials and housed for two days. This ensured that all focal females were 4-5 day old, sexually mature, and inseminated at the time of infections. Flies were again transferred into fresh food vials 8 hours before being subjected to experimental treatments (as described in the following subsection).

### Experimental design

Focal females were randomly distributed into three treatments: (a) sham-infected (n = 40 females per replicate), (b) infected with *E. faecalis* at OD600 = 0.8 (n = 80 females per replicate), and (c) infected with E. faecalis at OD600 = 1.5 (n = 80 females per replicate); the experiment was replicated thrice. After treatment, focal females were housed individually in food vials (with 8 mL food medium) for oviposition, and their survival was monitored every 1 hour for mortality for 48 hours. Previous experiments show that by this time point, infection-induced mortality plateaus in *E. faecalis*-infected females of this particular host population (Basu et al., 2024, Sehgal et al., 2025), and therefore, this window represents the acute phase of infection (*sensu* Howick and Lazzaro 2014).

For a subset of the dead females, randomly selected, the carcass was removed from the vial, and their systemic bacterial load was enumerated (see following subsection) within an hour of death; for the rest, their carcasses were discarded. This was done only for infected females, since no sham-infected female died in any replicate within the observation window. At 48 hours post-infection, observations were terminated, and 10 randomly selected living females from each treatment were used for the enumeration of systemic bacterial load (see following subsection); the rest of the living females were discarded.

The vials where the females had laid eggs were then incubated under standard maintenance conditions (see previous subsection) for the eggs to develop into adults; 12 days later, the number of adult progeny in each vial was counted. For each replicate of the experiment, extra females were infected (n = 30 females per infection dose per replicate) or sham-infected (n = 15 females per replicate) for the enumeration of bacterial load at 0 hours post-infection (see following subsection).

### Bacterial load measurements

Systemic pathogen load was enumerated for infected females (a) at 0 hours post-infection, (b) at death (i.e., bacterial load upon death, BLUD), and (c) at the end of the observation window (i.e., 48 hours post-infection).

For enumeration of bacterial load at 0 hours post-infection, females were surface sterilised twice (1 minute each) with 70% ethanol, washed in sterile distilled H_2_O once for 30 seconds, and homogenised in groups of 3 in 100 microliters of sterile MgSO_4_ (10 mM) buffer within 5-10 minutes of being infected. This homogenate was then serially diluted 1:10, eight times, in 100 microliters; 10 microliters of each dilution, along with 10 microliters of the original homogenate, was spotted onto Luria Bertani (Miller) agar (2%) plates. The plates were incubated for 8 hours at 37 ^O^C. Colony-forming units (CFUs) were counted for the countable dilution, and this count was multiplied by the appropriate dilution factor to calculate the systemic pathogen load. This was done for both infected (n = 10 pools of 3 females each per infection dose per replicate) and sham-infected (n = 5 pools of 3 females each per replicate) females; sham-infected female samples produced no CFUs in any of the replicates.

BLUD was enumerated for infected, dead females within 1 hour of their death. A protocol identical to the one above was used, except that the females were homogenised individually and in 200 microliters of sterile buffer. The sample size for each infection dose per replicate was 28-31 females.

For enumeration of bacterial load at 48 hours post-infection, again, an identical protocol was followed, except that the females were homogenised individually and in 50 microliters of sterile buffer. This was done for both infected (n = 10 females per infection dose per replicate) and sham-infected females (n = 10 females per infection dose per replicate); sham-infected female samples produced no CFUs in any of the replicates.

### Statistical analysis

All analyses were carried out using R statistical software, version 4.5.1 (R Core Team, 2025). Survival of infected females was analysed using mixed-effects Cox proportional hazards models with ‘infection dose’ as a fixed factor and ‘replicate’ as a random factor. Only data from infected females were included in the analyses, as sham-infected females did not exhibit any mortality within the observation window.

Female fecundity (progeny produced per hour) in each vial was normalised before analysis as follows:

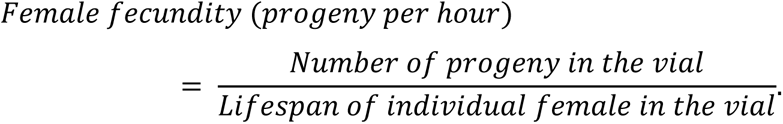

This was done to account for the fact that females differ in terms of how long they survive in the infected vs. the sham-infected treatments, and within the infected treatments, and therefore, there is a possibility that a female might produce a greater number of progeny by simply living longer than another female that perished early (Basu et al., 2024). This *normalised female fecundity* was analysed, using type III Analysis of Variance (ANOVA), separately with (a) ‘infection dose’ or (b) ‘infection outcome’ as fixed factors, and ‘replicate’ as a random factor.

0-hour bacterial load was analysed using type III ANOVA with ‘infection dose’ as a fixed factor and ‘replicate’ as a random factor. BLUD was analysed, using type III ANOVA, separately with (a) ‘infection dose’, (b) ‘time of death’ and ‘time of death × infection dose’ interaction, and (c) ‘female fecundity’ and ‘female fecundity × infection dose’ interaction as fixed factors, and ‘replicate’ as a random factors in every case. 48-hour bacterial load was analysed using type III ANOVA with ‘infection dose’ as a fixed factor and ‘replicate’ as a random factor.

## Results

### Post-infection survival

Infection with *Enterococcus faecalis* reduced survival of infected females compared to sham-infected controls (**figure 1(a)**); however, females infected with the higher infection dose (OD_600_ = 1.5) died equally compared to females infected at the lower infection dose (OD_600_ = 0.8; hazard ratio = 0.989, 95% confidence interval: 0.783-1.249, p = 0.928).

**Figure 1.**
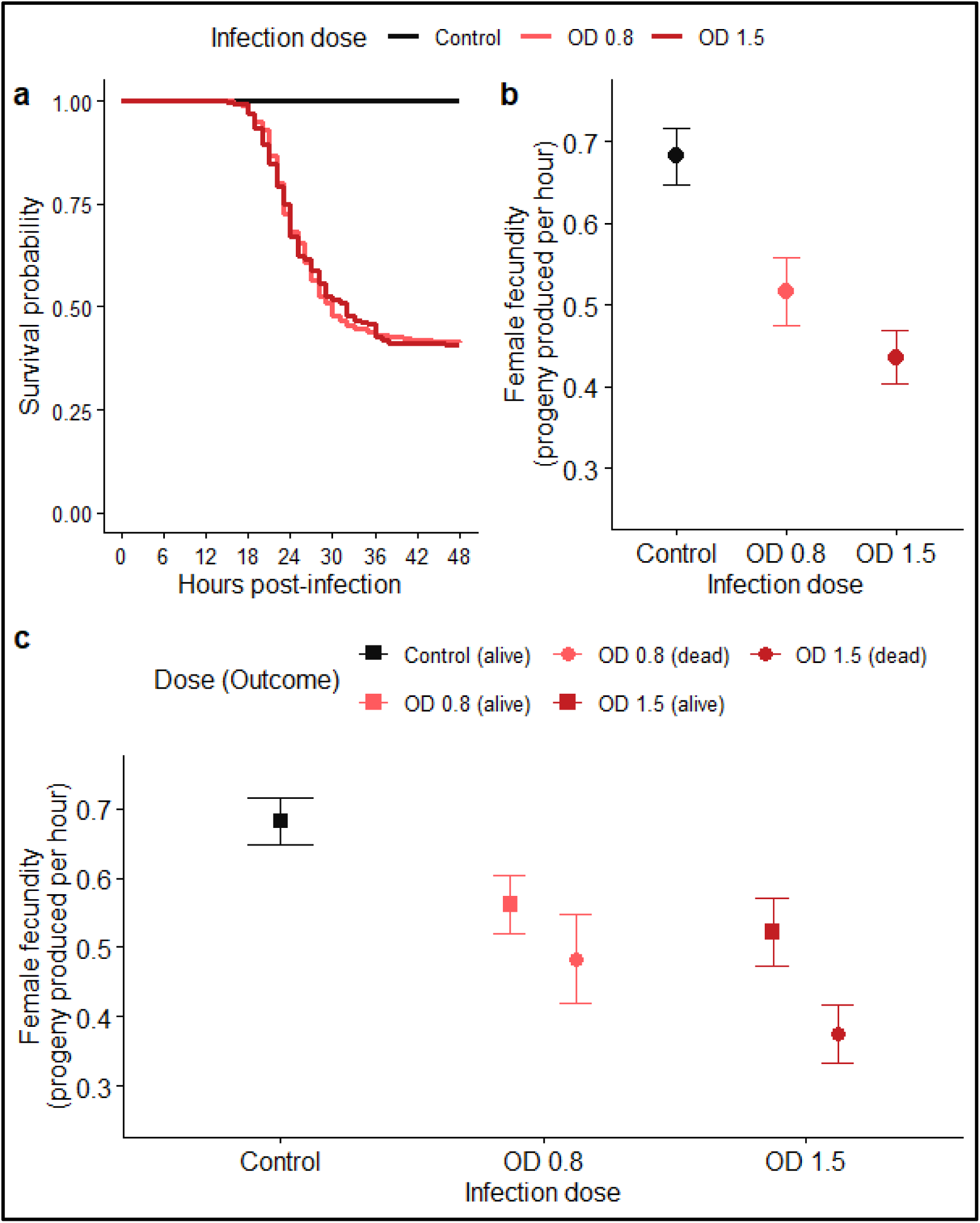
Effect of infection with *E. faecalis* on survival and fecundity of *D. melanogaster* females. **(a)** Effect of infection dose on survival; Kaplan-Meier plots. **(b)** Effect of infection dose on *normalised fecundity* (progeny produced per hour); mean ± 95% confidence interval. **(c)** Effect of infection outcome on *normalised fecundity*; mean ± 95% confidence interval.

### Post-infection fecundity

Infection with *E. faecalis* reduced fecundity of infected females compared to sham-infected controls in a dose-dependent manner (F_2,295_ = 32.579, p < 0.001; **figure 2(b)**; **supplementary table S1**). Females infected at the lower infection dose had significantly lower fecundity compared to controls (p < 0.001). Females infected at the higher infection dose had significantly lower fecundity compared to both controls (p < 0.001) and females infected at low dose (p = 0.001).

**Figure 2.**
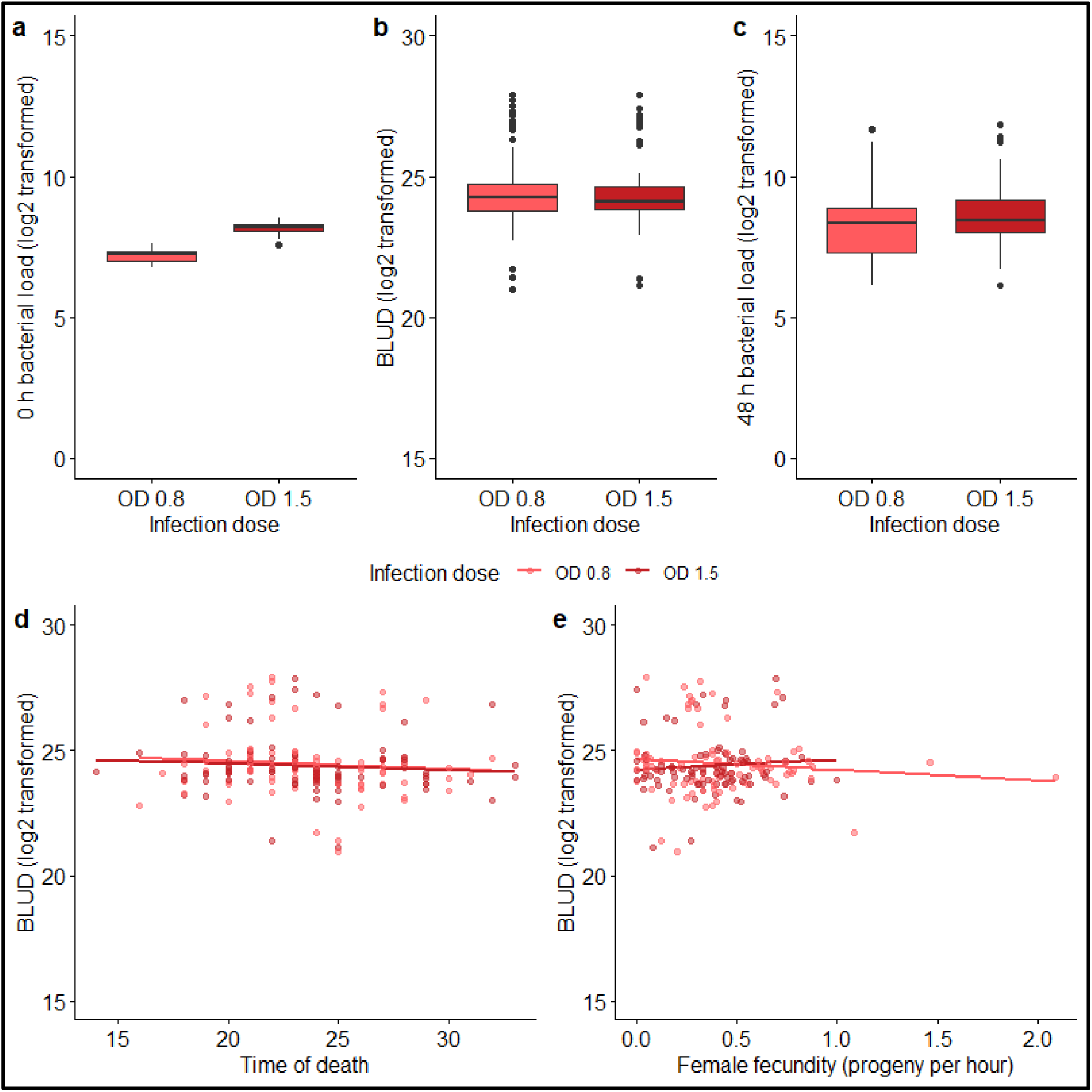
Bacterial load within females infected with *E. faecalis* and correlation with survival and fecundity. **(a)** Bacterial load at 0 hours post-infection. **(b)** Bacterial load upon death; BLUD. **(c)** Bacterial load (for survivors) at 48 hours post-infection. **(d)** Correlation between BLUD and time of death (hours post-infection). **(e)** Correlation between BLUD and female fecundity (progeny produced per hour).

Female fecundity was also contingent on infection outcome (F_4,597_ = 22.661, p < 0.001; **figure 2(c)**; **supplementary table S2**). For females infected at the lower dose, those who survived and those who died had significantly lower fecundity compared to controls (p = 0.011 and p < 0.001, respectively), but did not differ significantly from each other (p = 0.166). For females infected at the higher dose, those who survived and those who died had significantly lower fecundity compared to controls (p < 0.001 and p < 0.001, respectively), and also differed significantly from each other (p < 0.001), with those who died having the lowest fecundity. Across infection doses, females who were infected at high dose and survived had fecundity similar to those infected at low dose, whether they survived or died (p = 0.835 and p = 0.791, respectively). Females who were infected at high dose and died were significantly less fecund than those infected at low dose, whether they survived or died (p < 0.001 for both).

### Systemic bacterial loads

Females infected at the higher dose had significantly higher systemic bacterial load at 0 hours post-infection compared to females infected at the lower dose (F_1,60_ = 283.85, p < 0.001; **figure 2(a)**).

Infection dose (F_1,175_ = 0.256, p = 0.613) did not have an effect on bacterial load upon death (BLUD) of infected females (**figure 2(b)**). Time of death (F_1,79_ = 2.100, p = 0.151), or its interaction with infection dose (F_1,175_ = 0.2345, p = 0.629), did not have an effect on BLUD (**figure 2(d)**). Female fecundity (F_1,178_ = 0.320, p = 0.572), or its interaction with infection dose (F_1,176_ = 0.025, p = 0.875), did not have an effect on BLUD (**figure 2(e)**).

Infection dose did not have an effect on systemic bacterial loads of infected females at 48 hours post-infection (F_1,57_ = 0.445, p = 0.508; **figure 2(c)**).

## Discussion

We explored in this study the effects of infection dose (number of pathogen introduced in the host body at the point of infection) and infection outcome (whether a host dies or survives the infection) on host fecundity during the acute phase of infection, that is, when infection-induced mortality rates are high, and host fecundity is expected to be affected by the infection challenge (*sensu* Howick and Lazzaro 2014). To this end, we infected female *Drosophila melanogaster* flies with the pathogenic bacterium *Enterococcus faecalis* (Lazzaro et al., 2006) via septic injury at two different doses: OD_600_ of 0.8 and 1.5, *low-dose* and *high-dose*, respectively. Infection at *high-dose* introduced a greater number of bacterial colony-forming units into the fly body: females infected with the high dose had about 90% more CFUs in their system than those infected with the low dose, when systemic pathogen loads were enumerated at 0 hours post-infection (**figure 2(a)**).

Infection at *high-dose* did not significantly increase the risk of mortality: flies infected at either dose died at the same rate over the same period of time (**figure 1(a)**). An absence of an increase in mortality risk with increasing infection dose is surprising, but not entirely unexpected (Gupta and Vale 2017). On the other hand, increasing the infection dose had a significant effect on the infection-induced suppression of fecundity (**figure 2(b)**). While females infected with either infection dose produced significantly fewer progeny compared to control (sham-infected) females, the suppression of fecundity was greater in the case of females infected with the *high-dose*. Females infected with *high-dose* also produced significantly fewer progeny than females infected with *low-dose*. This suggests that the effect of the infection on host fecundity is independent of its effect on host mortality risk. A pathogen’s virulence — that is, the adverse effects of an infection challenge on host fitness — can manifest as reduced host survival and/or reduced host fecundity (Abbate et al., 2015). Our results thus demonstrate that, in female flies infected with *E. faecalis*, increasing the infection dose increases one aspect of the pathogen’s virulence without affecting the other. Suppression of fecundity following infection with *E. faecalis* has been previously demonstrated in other studies, too (Basu et al., 2024, Basu and Prasad 2024, Sehgal et al., 2025).

The suppression of fecundity in infected hosts is typically attributed to two mechanisms. It may either be caused by the diversion of limited resources from reproductive effort towards immune defences, including efforts to minimise damage (Sheldon and Verhulst 1996, Lochmiller and Deerenberg 2000, Schmid-Hempel 2005), or it may be a result of infection-induced damage to the host tissues and organs (Hurd 2001, Brandt and Schneider 2007). Since we observed that females infected at *high-dose* produced significantly fewer progeny than females infected at *low-dose*, one can suppose that this can be because either the hosts infected at high-dose invest a greater amount of resources into defences or that these hosts incur greater damage due to infection. We propose that some indication of which of these possibilities underlie our observations can be obtained by comparing the fecundity of females that survived the infection with that of those that did not across the two infection doses.

If surviving an infection at a higher infection dose requires greater resources, because the strength of immune response in the infected females is expected to positively correlate with infection dose (Tate and Graham 2017, Jent et al., 2019), then the two types of survivors would differ in progeny output, especially since both groups of survivors have successfully controlled the bacterial density within their bodies and limited it to comparable levels by 48 hours after infection (**figure 2(c)**). On the other hand, if the females that died of infection in the two infection treatments differ in progeny output, their differences may be attributed to differential systemic damage. Our results show that all infected survivors, irrespective of infection dose, exhibited similar progeny output (**figure 1(c)**). However, amongst the females that died, the females infected at *high-dose* produced significantly fewer progeny compared to the ones infected at *low-dose* (**figure 1(c)**). This suggests that infection-induced damage is more likely to explain our results across the two infection doses. Our assertion is supported by previous work showing that, within the same experiment, infection with *E. faecalis* suppressed female fecundity, but altering reproductive effort did not affect post-infection survival, suggesting no involvement of resource-allocation trade-offs (Sehgal et al., 2025). Furthermore, *high-dose* infected-dead females produced significantly fewer progenies compared to females in all other dose × outcome combinations, all of whom had similar progeny output (**figure 1(c)**).

Why does an infected host die? It can be supposed that the total accumulated damage — caused by the pathogen and the host’s own immune response — to the host soma kills an infected fly (Shirasu-Hiza and Schneider 2007, Duneau et al., 2017, Duneau et al., 2025). A potential proxy for this damage can be the bacterial load carried by infected flies at the time of their deaths, formally termed bacterial load upon death, or BLUD (Duneau and Ferdy 2022). BLUD for a pair of host and pathogen strains shows little variance, and can thus be used as a proxy for host tolerance (Duneau et al., 2017), or, in other words, how much damage a host can sustain before it succumbs to the infection (Duneau and Ferdy 2022, Huang et al., 2023). Immunopathology, stemming from the host’s immune response, can also affect BLUD (Duneau et al., 2025). We had therefore also measured BLUD of the females of both infection treatments in our experiment.

Our results show the BLUD is not affected by the infection dose: females of both infection treatment carried similar individual bacterial loads at the time of their death (**figure 2(b)**). Additionally, BLUD was unaffected by the time of death of the females (**figure 2(d)**). Existing literature suggests that independence from both infection dose and time of death are characteristic properties of BLUD (Duneau et al., 2017, Jent et al., 2019, Wilson et al., 2020, Vincent et al., 2020), and therefore, our results agree with the set precedent. However, if BLUD is taken as a proxy of damage incurred, our results also indicate that females in both infection treatments incurred comparable levels of damage. But, again, we observed that these females produced different number of progeny. We further tested and found that even at the level of individual females, BLUD was not correlated with the females’ progeny output (**figure 2(e)**).

We therefore argue that BLUD might not serve as a suitable proxy for total damage incurred by the host, but only for the contributions of the pathogen towards this total. We additionally argue that the somatic damage that kills the host (for which BLUD serves as a good proxy) is independent of the damage to reproductive tissue (which leads to fecundity suppression). This, if true, will explain why fecundity suppression intensifies with increasing infection dose in our experiments without a concomitant increase in mortality risk. Furthermore, previous work in *D. melanogaster* has shown that infection combined with damage to only particular tissues increases mortality risk (Chambers et al., 2014), which supports our assertion.

In summary, we have demonstrated in this study that increasing the infection dose of the bacterial pathogen *E. faecalis* can lead to greater suppression of female fecundity in *D. melanogaster*, without necessitating a concomitant increase in host mortality risk. We also find that hosts that die from infection can have a different reproductive output compared to the hosts that survive the infection challenge. However, this difference might only be observable at specific infection doses and not others. We propose that greater fecundity suppression following infection at a higher dose may result from hosts incurring greater systemic damage, particularly to their reproductive organs. We hope that further studies will elucidate this phenomenon in greater detail. Furthermore, our results may be unique to our study population or to the pathogen strain we used. Hence, the generalizability of our results to other host-pathogen systems is contingent on independent replication of our results. To conclude, a variety of host and pathogen features, and infection parameters, are known in the extant literature to determine the influence of bacterial infection on fly fecundity. Our study adds two more factors — infection dose and infection outcome — to that list.

## Author contributions (CRediT statement)

**Aabeer Kumar Basu** (Conceptualisation, Methodology, Investigation, Data curation, Validation, Formal analysis, Visualisation, Writing – original draft, Writing – review & editing), **Kimaya Tekade** (Methodology, Investigation, Data curation, Writing – review & editing), and **Nagaraj Guru Prasad** (Funding acquisition, Supervision, Writing – review &editing)

## Declaration of generative AI tools

No generative AI tools were utilised during the writing of this manuscript or during any other process concerning the experiments reported in this manuscript.

## Acknowledgements

The authors thank Prof. Brian Lazzaro (Cornell University, USA) for providing the *Enterococcus faecalis* isolate used in the experiments.

