## Supplementary Tables 1 and 2 for "Dose- and outcome-dependent effects of bacterial infection on female fecundity in *Drosophila melanogaster*"

**Supplementary table S1.** Effect of infection dose on female fecundity: post hoc Tukeys HSD for pairwise comparisons.

| **(a) Least-square means.** | | | |
| --- | --- | --- | --- |
| Treatment/infection dose | Least-square mean | Lower CI | Upper CI |
| Sham-infected controls | 0.682 | 0.623 | 0.741 |
| Low dose (OD_600_ = 0.8) | 0.516 | 0.466 | 0.565 |
| High dose (OD_600_ = 1.5) | 0.435 | 0.386 | 0.485 |

| **(b) Pairwise comparisons.** | | | | | |
| --- | --- | --- | --- | --- | --- |
| Comparison | Estimate | SE | DF | t ratio | p value |
| Sham – low dose | 0.167 | 0.031 | 597 | 5.432 | **< 0.001** |
| Sham – high dose | 0.247 | 0.031 | 597 | 8.060 | **< 0.001** |
| Low dose – high dose | 0.080 | 0.031 | 597 | 3.202 | **0.004** |

**Supplementary table S2.** Effect of infection outcome on female fecundity: post hoc Tukeys HSD for pairwise comparisons.

| **(a) Least-square means.** | | | | |
| --- | --- | --- | --- | --- |
| Treatment/infection dose | Infection outcome | Least-square mean | Lower CI | Upper CI |
| Sham-infected controls | na | 0.682 | 0.622 | 0.742 |
| Low dose (OD_600_ = 0.8) | Survival | 0.563 | 0.499 | 0.627 |
| Low dose (OD_600_ = 0.8) | Death | 0.482 | 0.425 | 0.540 |
| High dose (OD_600_ = 1.5) | Survival | 0.523 | 0.459 | 0.586 |
| High dose (OD_600_ = 1.5) | Death | 0.375 | 0.318 | 0.432 |

| **(b) Pairwise comparisons.** | | | | | |
| --- | --- | --- | --- | --- | --- |
| Comparison | Estimate | SE | DF | t ratio | p value |
| Sham – low/survival | 0.119 | 0.037 | 602 | 3.237 | **0.011** |
| Sham – low/death | 0.200 | 0.034 | 601 | 5.949 | **< 0.001** |
| Sham – high/survival | 0.159 | 0.037 | 600 | 4.344 | **< 0.001** |
| Sham – high/death | 0.307 | 0.033 | 600 | 9.193 | **< 0.001** |
| Low/survival – low/death | 0.080 | 0.036 | 603 | 2.241 | 0.166 |
| Low/survival – high/survival | 0.040 | 0.038 | 600 | 1.044 | 0.835 |
| Low/survival – high/death | 0.188 | 0.036 | 603 | 5.267 | **< 0.001** |
| Low/death – high/survival | -0.040 | 0.036 | 603 | -1.129 | 0.791 |
| Low/death – high/death | 0.107 | 0.032 | 599 | 3.348 | **0.007** |
| High/survival – high/death | 0.148 | 0.035 | 602 | 4.162 | **< 0.001** |
